# Navigation of Ultrasound-controlled Swarmbots under Physiological Flow Conditions

**DOI:** 10.1101/2022.02.11.480088

**Authors:** Alexia D.C. Fonseca, Tobias Kohler, Daniel Ahmed

**Author notes:** Correspondence and requests for materials should be addressed to D.A.

## Abstract

Navigation of microrobots in living vasculatures is essential in realizing targeted drug delivery and advancing non-invasive surgeries. We developed acoustically-controlled “swarmbots” based on the self-assembly of clinically-approved microbubbles. Ultrasound is noninvasive, penetrates deeply into the human body, and is well-developed in clinical settings. Our propulsion strategy relies in two forces: the primary radiation force and secondary Bjerknes force. Upon ultrasound activation, the microbubbles self-assemble into microswarms, which migrate towards and anchor at the containing vessel’s wall. A second transducer, which produces an acoustic field parallel to the channel, propels the swarms along the wall. We demonstrated cross- and upstream navigation of the swarmbots at 3.27 mm/s and 0.53 mm/s, respectively, against physiologically-relevant flow rates of 4.2 – 16.7 cm/s. Additionally, we showed swarm controlled manipulation within mice blood and under pulsatile flow conditions of 100 beats per minute. This capability represents a much-needed pathway for advancing preclinical research.

**Teaser:** Navigation of ultrasound-guided microrobots inside artificial blood vessels overcoming physiological conditions, including high flow rates, pulsatile flow regimes, and high cell concentrations of blood.

## Introduction

Navigation of miniaturized robots in vasculature under physiological flow conditions could become an exciting prospect for treating numerous diseases. For example, replacing conventional systemic treatments with targeted approaches that deliver doses locally to a lesion promises to improve therapeutic efficacy for an array of vascular diseases such as stroke, peripheral artery disease, abdominal aortic aneurysm, carotid artery disease, pulmonary embolism (blood clots), deep vein thrombosis, atherosclerosis, and cancer. A key drawback of conventional systemic treatments is that the initial drug concentration becomes diluted across the total circulating blood volume, thus significantly reducing its efficacy. Beyond increasing drug effectiveness, targeted therapy also minimizes drug side effects by avoiding interference with healthy organs. However, despite the great potential of microrobots in many areas of medical research, their manipulation inside the vasculatures of animal models still poses a fundamental challenge and there remain numerous limitations to be overcome(*1*). Physiological conditions inside living systems involve high flow velocities, pulsatile flow regimes in large arteries, and the need to coexist with cellular components. All in all, microrobots must confront an extremely challenging environment and be carefully guided through a patient’s vasculature. To date, efforts in this field have focused on controlling microrobot trajectory inside microchannels and confined spaces(*2*–*8*), and most studies have aimed to manipulate microrobots under static flow conditions(*9*–*12*). Recent work has begun investigating microrobot capabilities when exposed to an external flow(*13*–*16*), but typically considers only low flow velocities, i.e., in the μm/s range. When exposed to physiological flow values that are in the cm/s range, as found in mice and humans, most engineered microrobots cannot withstand the drag force and are swept away.

Developing synthetic systems that can propel upstream would significantly increase the scope of *in vivo* microrobot applications. Recent research has tended to exploit the motion of synthetic swimmers near boundaries, where non-slip conditions minimize flow velocity, drag force is reduced, and upstream manipulation is easier. Catalytic approaches have been widely investigated(*17*–*22*), and Katuri et al. demonstrated a controlled cross-stream motion of janus particles, which could mimic the motion of natural microswimmers and be exploited to reach vessel boundaries(*23*). Importantly, some approaches have already managed to manipulate microswimmers upstream when they are close to a wall. Ahmed et al. exploited a rotating magnetic field combined with ultrasounds to roll microrobots upstream along a vessel wall(*24*, *25*). Additionally, Alapan et al. combined magnetically-guided rolling upstream manipulation with antibody tumour targeting(*26*). In a third example, Wang et al. used magnetic swarms that could assemble in stagnant conditions and subsequently be manipulated upstream(*27*). However, while these recent approaches all permit microrobots to execute upstream motion, they still exhibit slow velocities and microrobot control against flow. Furthermore, magnetic actuation platforms(*28*–*34*) predominate, but face difficulty when scaling up and reproducing the behaviour in more complex environments. Till date, we lack effective ultrasound-based microrobots performing upstream propulsion in physiological flow conditions(*28*).

Ultrasound in particular is an attractive modality for navigation of micro/nanorobots *in vivo* because it penetrates deep into the tissue, is not affected by the opaque nature of animal bodies, is relatively non-invasive, and generates a broad range of forces. More importantly, ultrasound imaging systems are already widely used in clinical settings for the real-time imaging of objects, and such systems have recently been utilized for tracking microrobots(*27*, *35*, *36*). Ultrasound-based microrobots are gradually becoming popular because the designs are relatively simple, their fabrication is inexpensive and does not require doping with magnetic particles, their experimental setup is likewise inexpensive, and their operation is simple(*37*–*41*). Resonating microbubbles confined in soft microfabricated shells that are capable of extremely fast and selective micro-propulsion have been utilized in 3D manipulation(*42*–*44*). However, while resonating microbubbles confined in cylindrical shells generate large propulsive forces, the microbubbles are typically not stable, i.e., they grow over time; as such, driving them becomes a fundamental challenge. Researchers have also investigated the self-assembly of microbubbles(*45*–*47*), and to date, most such studies involve only trapping of microbubbles or microparticles when exposed to an external flow(*48*, *49*). Despite the interest in and potential of acoustic-based microrobots of all types, a thorough investigation of how they navigate when exposed to physiological flow rates has yet to be made.

In this article, we introduce acoustic swarmbots capable of circulating through the vasculature, aiming to overcome the aspects of physiological environments that limit external microrobot control (Fig. 1A). The swarm design we propose relies purely on acoustic forces for self-assembly, and those forces are introduced from two perpendicular transducers, facilitating an easy setup. First, activating one piezoelectric transducer induces an acoustic wavefield and associated forces that cause the microbubbles to self-organize into a microswarm. Secondly, the primary acoustic radiation force pushes the microswarm away from the acoustic transducer and towards the wall of the blood vessel (Fig. 1B). Upon the MBs reaching the wall, the second piezoelectric transducer is activated, the radiation force from which enables upstream propulsion (Fig. 1B,C). Importantly, vessels inside an organism present laminar flow, which features zero velocity at the wall. Our system exploits this intrinsic characteristic: the simultaneous assembly and migration towards the wall allows our swarmbots to overcome high flow rates and avoid the usual limitations that hinder manipulation; our microrobots are able to move upstream with velocities of up to 0.53 mm/s against a flow of 4.2 cm/s. Furthermore, the swarm feature of the microrobots enables them to withstand physiological conditions, including blood flow velocities of up to 16 cm/s, over any desired range of time. Finally, our microswarms have the potential to be controllable in any blood vessel, from the smallest capillaries to the largest arteries; their collective behaviour allows the robots to adapt to vessel size and shape, on top of being able to navigate within a pulsatile flow regime. This technology has great potential for impact as a foundation for advancements in brain research and vasculature biology, as well as prompting the development of relevant targeted treatments.

**Fig. 1.**
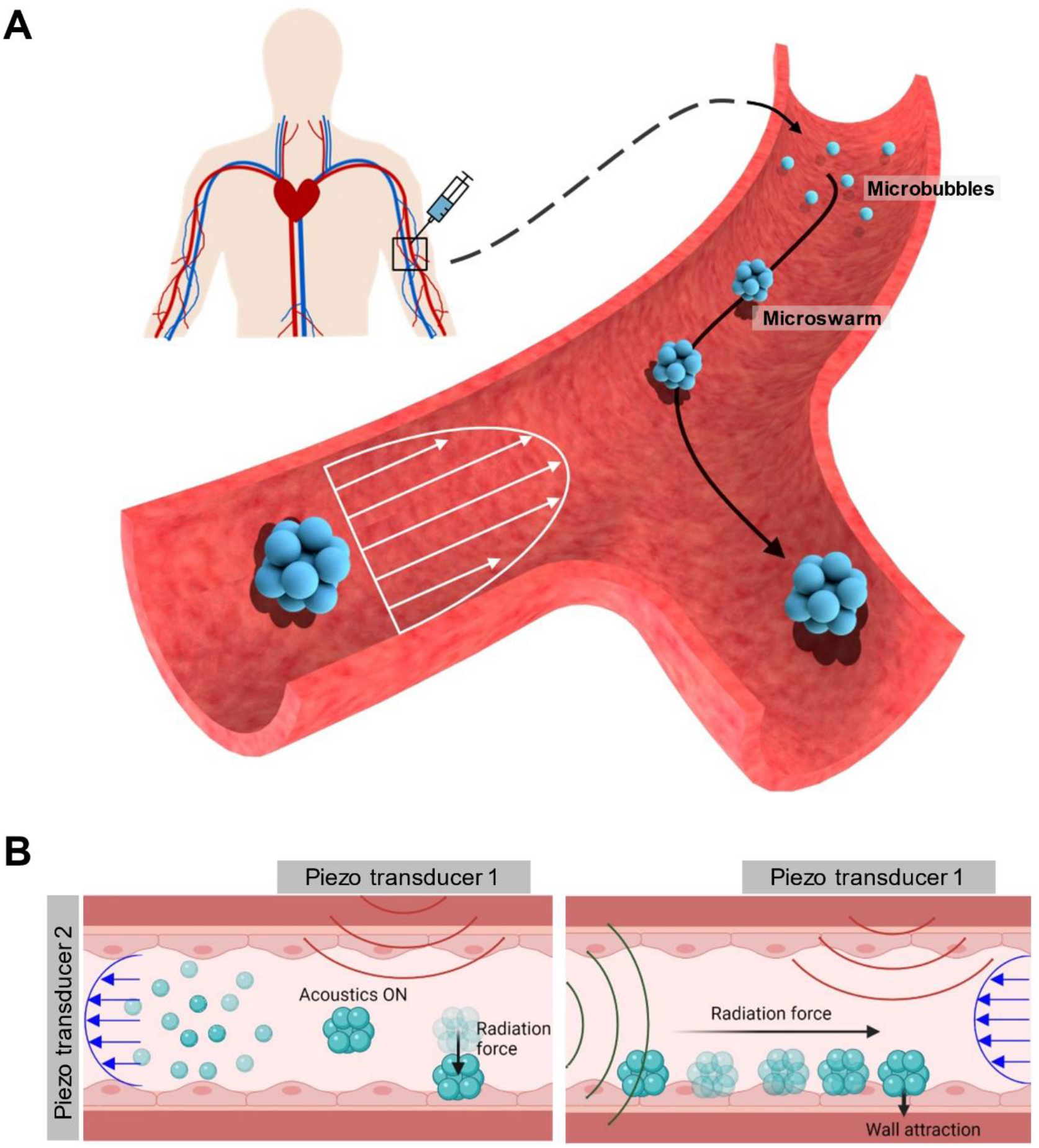
Concept of ultrasound-controlled swarmbots. **(A).** Saline solution containing dispersed MBs is used to introduce the microrobots into the patient via injection. When excited with ultrasound, the single microbubbles self-assemble into a swarm. These microswarms can be manipulated upstream and cross-stream using only acoustic actuation. **(B).** Self-assembly and manipulation of microrobots is performed using two piezoelectric transducers in combination. One actuator is placed parallel to the vessel (piezo transducer 1); when activated, the microswarm assembles and migrates towards the vessel wall. The second actuator is placed perpendicular to the channel (piezo transducer 2); its activation leads to microswarm propulsion away from the transducer.

## Results

### Experimental setup

The working principles of our microrobots were analysed in microfluidic channels to elucidate the fundamental mechanisms underlying acoustic microswarm propulsion. Our acoustic chamber comprised a transparent polydimethylsiloxane (PDMS)-based microfluidic device with a square cross-section of 400 × 30 μm. We chose PDMS because its acoustic properties are similar to those of biomaterials that exist in the human body, such as blood, fat, the brain and other soft tissues. In addition, the transparent nature of PDMS allows for excellent visualization by brightfield and fluorescent microscopy. The acoustofluidic device also featured a pair of piezoelectric transducers, PZT 1 and PZT 2, which were positioned orthogonal to each other (Fig. S1). The piezoelectric transducers were bonded on the chamber’s sidewalls, ensuring that travelling acoustic waves predominated, and were connected to a pair of electronic function generators that regulated the frequency and amplitude of the produced ultrasound. The transducer pair was activated at frequencies close to their resonance, i.e., between 240 – 250 kHz, and with peak-to-peak voltages of 0.1 – 20 V_PP_. The frequency spectrum of the transducers is given in Fig. S2. Finally, we injected commercially-available, biocompatible, 1 – 2-micron gas-filled polymeric-shelled microbubbles (Sonovue, Bracco Imaging) through a programmable syringe pump. These microbubbles (MBs) consist of a thin lipid monolayer filled with inert sulphur hexafluoride gas and remain stable for hours. The whole setup was positioned on an inverted microscope, and experimental data were recorded using high-speed and high-sensitivity cameras.

### Mechanism of acoustic microswarm formation and its translational motion

In this section, we combine multiple physical principles of acoustic MBs in the setting of artificial vessels and investigate their application as a microrobotic navigation platform. We used two piezoelectric actuators to excite the MBs (Fig. 1B): Piezo transducer 1, positioned parallel to the artificial vessel, which induced self-assembly and migration towards the wall; and Piezo transducer 2, placed perpendicular to the vessel, which controlled propulsion along the wall. We specifically studied the self-assembly and manipulation of the acoustically activated MBs.

We injected gas-filled MBs into the artificial vessel, where they were immersed in liquid. When these MBs were exposed to ultrasound, they oscillated and became excellent scatterers of the sound field. This effect can be attributed to the large acoustic impedance mismatch that exists at a liquid/air interface, where acoustic impedance measures the reflection and transmission of a sound wave at a boundary. When acoustic waves propagating through the liquid environment encounter the gas medium within the MBs, the liquid/air interface strongly reflects the energy of the incident waves. Cross-scattering of these reflections from multiple MBs generates a localized pressure gradient and gives rise to secondary radiation forces, F_B_, which act on adjacent MBs. For compressible MBs in a water solution, these forces can be described by the following relation, assuming the driving force is much less than the resonance frequency of the microbubbles:

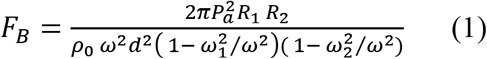

where *ω* and *P_a_* are the excitational ultrasound frequency and acoustic pressure, respectively; *R_1_* and *R_2_* are the radii of the MBs, *p_0_* is the density of the liquid; *d* is the distance between bubble centers and *ω*_1_ and *ω*_2_ are the resonance frequencies of the bubbles. Typically, the mutual force between two bubbles is attractive when they oscillate in phase and repulsive when out of phase. Here, since the neighbouring MBs are approximately equal in size and were activated by the same frequency, they oscillate in phase; thus, upon ultrasound excitation by Piezo transducer 1 at 242 kHz and 1 V, the MBs were instantaneously attracted, which initiated the self-assembly process (Fig. 2A). When stabilized, which required only milliseconds, this self-assembly resulted in highly ordered circular spheres composed of 100 to 200 MBs (Fig. 2B & Movie S1). The assembly process thus qualitatively agrees with Eq. 1. We further characterized the self-assembly process across a range of MB concentrations, ultrasound frequencies, and voltages.

**Fig. 2.**
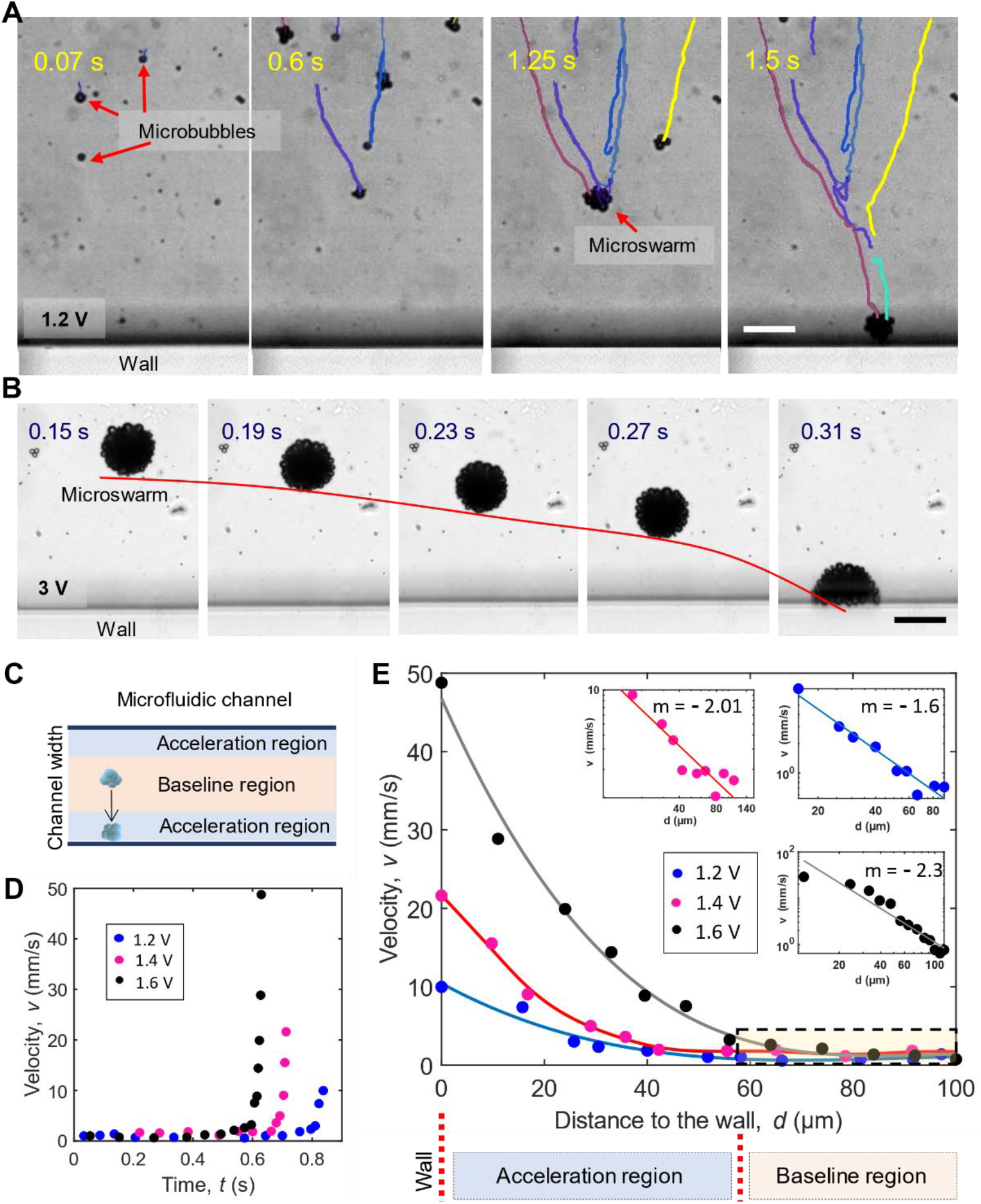
Self-assembly and migration of microswarms towards a channel wall. **(A).** Injected microbubbles under an acoustic excitation field of 1.2 V and 242 kHz from a piezo transducer positioned parallel to one channel wall simultaneously self-assemble and migrate towards the opposite wall. Scale bar, 50 μm. **(B).** Image sequence demonstrates migration of a self-assembled microswarm towards the wall under an acoustic excitation field of 3 V and 242 kHz. The trajectory, marked in red, shows an acceleration when the microswarm approaches the wall. Scale bar, 50 μm. **(C).** Two distinct regions are identified inside the channel—a core region where microrobot velocity is nearly constant, and an acceleration region bordering each wall, where microbubbles accelerate towards the wall. **(D).** Plot showing microswarm velocity over time during migration towards the wall. **(E).** Plot demonstrating the relationship of microswarm migration velocity to distance to the wall. Distances corresponding to baseline and acceleration regions are indicated by yellow and blue boxes, respectively. Logarithm analyses are shown for the three curves plotted.

As the MBs continuously self-assembled, they simultaneously were migrated towards the wall, (Fig. 2A & Movie S2). We observed that these microswarms were pushed away from Piezo transducer 1 and become trapped into the PDMS wall (Fig. 2B & Movie S3). The navigation of the microswarms can be attributed to the gradient of the acoustic field generated by the incident pressure field from Piezo transducer 1. This gradient exerts a primary radiation force, which can be expressed by the following relation:

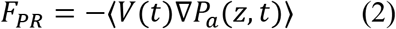

where 〈 〉 is the time average, *V*(*t*) is the instantaneous volume of the bubble and *P_a_*(*z*, *t*) is the external acoustic pressure acting on it. The force generated from Piezo transducer 1 introduced a ‘pushing’ action on the MBs, i.e., the MBs were driven towards the opposite vessel wall. Once the MBs reached the wall, we activated Piezo transducer 2 such that microswarms were pushed forward along the vasculature wall by again concept primary radiation force but now acting perpendicular to the force generated by Piezo transducer 1 (Movie S4). The propulsion direction can be tuned, i.e., with or against the flow, by positioning the piezo actuator facing or opposite the flow direction. Although microswarm formation and migration towards the wall are consistent with previous findings(*45*–*47*), we devise a strategy that apply these phenomena in developing an efficient navigation platform that can execute cross-stream and upstream motion against physiological flow conditions.

We aim to exploit the non-slip boundary condition at walls of the artificial vessels, where the flow velocity tends to be zero, thus, facilitating efficient microrobot manipulation. For this, we characterized microswarm behaviour when moving towards the wall. Interestingly, as the acoustic microswarms approach the wall, they start to accelerate, i.e. there is a transition point during microswarm migration at which swarm velocity starts increasing (Fig 2D). This transition can be attributed to secondary Bjerknes forces. Typically, when two MBs of equal size oscillating in an acoustic field are placed close to one another, they are mutually attracted by the secondary Bjerknes force. Similarly, when an oscillating MB is placed near a wall or surface, it is attracted to the wall. This attractive force can be understood by placing an imaginary microbubble on the opposite side of the wall. When real bubble 1 oscillates at a particular phase, mirror microbubble 2 oscillates at the same phase; thus, the real bubble is attracted to the wall. The velocity of a MB approaching a wall due to secondary radiation forces can be derived equating Eq. (1), where we assume *R*_1_ = *R*_2_ = *R*_0_ and *ω_1_* = *ω_2_* = *ω_0_*, to the Stokes drag force: *F_d_* = –*CπηR_0_v* Eq. (3), where *v* is the velocity of MBs. Thus, it can be written as 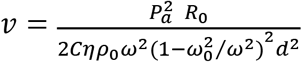 Eq. (4), where for *Re ≪ 1, C* = *4*, *Re* denoting the Reynolds number, *η* is the viscosity term, and *C* is the drag coefficient. Note that the Reynolds number of microbubble navigation can be estimated as 0.043. As microswarm velocity scales inversely with a power of 2 in relation to swarm distance to the wall (Fig. 2E), our experimental data coincide with Eq. (4).

Based on microswarm behaviour in response to excitation by Piezo transducer 1, we determined there to be two distinct regions within a vessel (Fig. 2C). At the centre of the vessel, where contribution from the wall is negligible, MBs tend to move with an almost uniform velocity; we term this the ‘baseline region.’ At a certain distance from the wall, the MBs sense the wall and initiate acceleration, hence we term this area the ‘acceleration region’. Mechanistically, the secondary (Bjerknes) radiation force increases as distance from the wall decreases; thus, secondary forces become sufficiently large so as to trap the microswarm at the wall. The transition that distinguishes the two regions is not a constant, i.e., the size of the acceleration region varies with voltage, frequency, and MB concentration. All told, we have characterized microswarm manipulation towards the wall, and trapping at the wall upon ultrasound excitation of Piezo transducer 1. Understanding these events is the first step towards implementing microswarm propulsion along a wall. Piezo transducer 2 will play key role in this manipulation, as shown in Fig.3; tuning the position of this transducer we will move microswarms up or down stream inside the vessels.

**Fig. 3.**
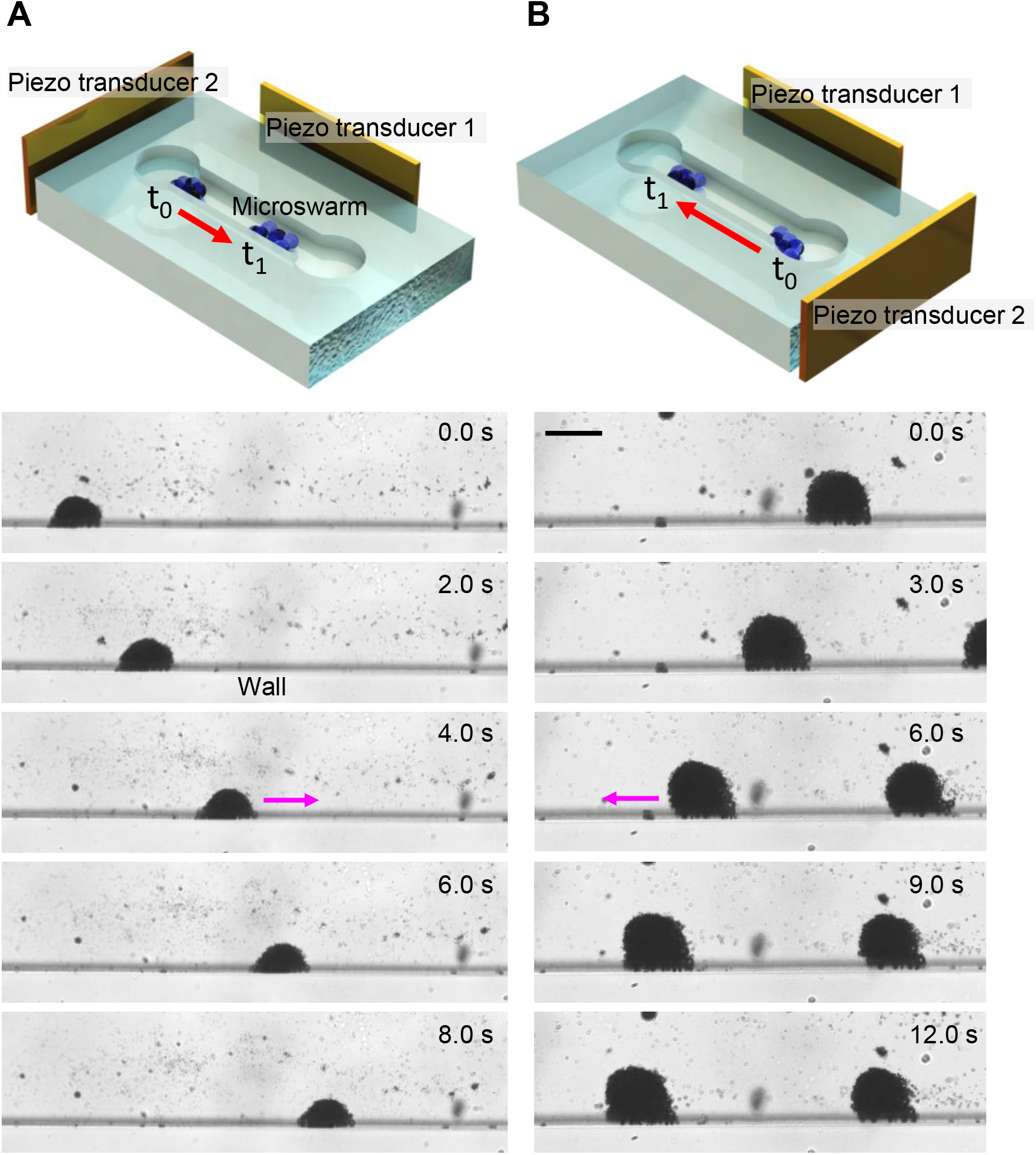
Ultrasound-controlled microswarm navigation in static solution. **(A)**. A 3D model of the experimental setup showing the positions of piezo transducers 1 and 2. Piezo transducer 2 is placed at the left, perpendicular to the vessel. When it is activated, the swarm is pushed towards the right, sliding along the vessel wall. **(B).** When piezo transducer 2 is placed on the opposite side of the channel, i.e. the right, the swarms are subsequently pushed towards the left. Scale bar, 50 μm.

### Controlling swarming behaviour: swarm size and assembly dynamics

We performed a parametric study to elucidate factors that govern microswarm size and the rate at which swarms are produced. This necessitated studying two different phases of MB behaviour: the initial stage when individual MBs start moving towards each other and towards the wall, termed ‘single bubble motion’; and the movement of swarms after having formed and reached a specific diameter, termed ‘swarm motion’. These different phases are illustrated in Fig. S2A.

This section concerns the ‘single bubble motion’ phase during which acoustic microswarms are created, and specifically the effects of voltage, frequency, and MB concentration on swarm formation. Notably, self-assembly occurs throughout the vessel, but those clusters that assemble close to a wall undergo simultaneous acceleration towards the wall. To analyse this self-assembly independently from wall attraction, we specifically assessed swarm behaviour within the ‘baseline region’ of the vessel, at least 60 ± 10 μm from both vessel walls. Speed in Fig. S2 correspond to ‘baseline region’, v_B_, that is, the speeds of MBs on their way to swarm assembly. As described in Fig. S2B, the velocity of a single MB increased from 5 to 100 μm/s when the voltage applied to the piezo transducer increased from 0.2 to 0.9 V_PP_. Thus, increasing the voltage or acoustic pressure by 4.5 times causes MB velocity to increase by 1900%. These experimental data match the relationship between pressure and velocity described in Eq. (4), as velocity scales with a power of two when increasing the voltage applied to the transducer.

Additionally, we studied the dependency of individual MB velocity on acoustic frequency across the range of 110 kHz to 300 kHz, i.e. values close to the transducer resonance frequency of 244.8 kHz; see Fig. S2D. A maximum velocity of 60.88 ± 19.21 μm/s was produced at ~220 kHz. We next investigated the effect of MB concentration, testing 2, 6, 8, 10, 15, and 20 MB μl/ml in saline solution at 242 kHz and 0.6 V_PP_, and achieved a maximum velocity of 98.01 ± 33.21 μm/s at 10 μl/ml. Thus, we determined optimal values for frequency (~220 kHz) and MB concentration (~10 μl/ml), as shown in Fig. S2C,D. We also identified that the effect of MB concentration on velocity is greater than that of frequency. However, increasing MB concentration by five times caused a velocity increase of up to ~440%, while increasing frequency by 1.3 times yielded a velocity increase of as much as ~6000%. Consequently, the system is more sensitive to frequency changes. Note that the optimal frequency value (~220 kHz) nearly coincides with the resonant frequency of the piezo transducer (244.8 kHz, Fig. S2D); accordingly, the optimal frequency can vary between different piezoelectric actuators. Overall, we can regulate the rate of microswarm self-assembly by selecting an appropriate voltage, frequency value, and MB concentration.

We then studied the parameters controlling swarm size upon activation of the acoustic signal. We first investigated the effect of MB concentration. When maintaining constant power and frequency, increasing MB concentration by three times caused microswarm size to increase 105%. When using the aforementioned optimal MB concentration of ~10 μm/ml, microswarms reached a saturation diameter at ~120 ± 5 μm (Fig. 4A). Inclusion of time as a fourth variable enabled control of cluster size via voltage. When we increased the excitation voltage from 0 to 3 V_PP_, single MBs migrated at higher speeds, hence larger microswarms were produced rapidly (Fig. 4B,C). When using voltages between 1 and 3 V_PP_, microswarms were assembled with diameters of 85 ± 65 μm (Fig. 4B). Here, we elucidated various parameters regulating microswarm formation (speed and swarm size), establishing the foundation for implementing a controlled microrobotic approach. In the following sections, we focus on translating this approach in physiological flow conditions.

**Fig. 4.**
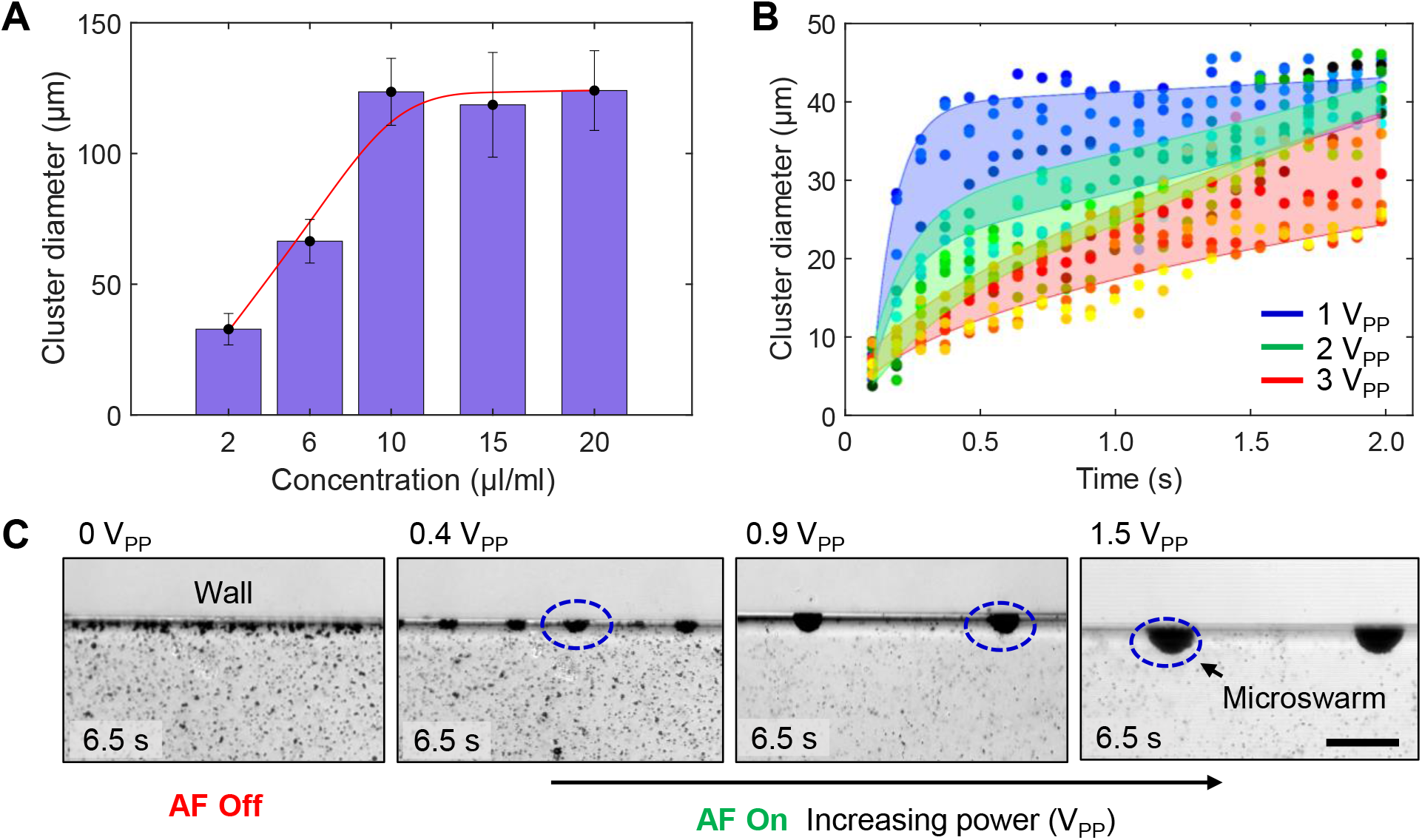
Characterization of microswarm size. **(A).** Bar plot of microswarm cluster diameters at different initial MB concentrations. Each bar represents the average cluster diameter determined from five microrobots. Error bars represent standard deviation (s.d.). **(B).** Plot demonstrating the time evolution of cluster diameter in five different microswarms at 1, 2, and 3 V_PP_. **(C).** Image sequences demonstrating microswarm size at 0, 0.4, 0.9, and 1.5 V_PP_ over a uniform 6.5-second period. Scale bar, 50 μm.

### Microswarm manipulation in flow conditions

Under physiological conditions in living systems, such as in humans, the preeminent obstacle is high blood flow velocities in the cm/s range. Moreover, the human circulatory network includes vasculature with varied cross-sectional diameter and flow profile. In large arteries, the pumping heart generates a pressure gradient that produces high flow velocities and a pulsatile profile; previous studies have documented flow velocities of 5 – 19 cm/s in human arteries(*51*), with big vessels such as the aorta potentially reaching peak values of 66 cm/s(*52*). In addition, the arteries are split and divided into progressively smaller branches, ultimately transitioning into capillary networks that irrigate organs with nutrients and oxygen. As vessel diameter decreases, its resistance, i.e., the force opposing fluid flow, increases, hence lower and non-pulsatile flow velocities exist within capillaries. By the time blood has passed through the capillaries and reached the veins, the pressure gradient is almost negligible and pulsatile flow has been damped, but still blood flow velocities range 1.5 – 7.1 cm/s(*51*); with big veins such as the vena cava can reach peak values of 28 cm/s(*52*). Here, to test the translatability of our approach to *in vivo* applications, we implemented conditions as close to the physiological flow in humans as possible. We have considered a wide range of possible conditions, including a range of flow velocities and possible pulsatile flow regimes. For the purpose of this assessment, we classified microswarm motion as: cross-stream (v_cross_), which is perpendicular to the stream direction; upstream (V_up_), which is against the stream direction; and downstream (V_down_), which follows the direction of the flow (Movie S5).

We first characterized the cross-stream motion during and after self-assembly that resulted from Piezo transducer 1 actuation. That is, when acoustically activated in the absence of any flow, the microswarms transversed perpendicularly to the vessel wall, a motion attributed to a combination of primary and secondary radiation forces. When exposed to an external flow, swarms moved towards the wall at an angle due to the combination of acoustic force from Piezo transducer 1 and downstream drag forces. In addition, larger microswarms were ‘pushed’ towards the wall more strongly, i.e., they exhibited trajectories more perpendicular to the stream, while smaller microswarms demonstrated more parabolic movement (Fig. 5A). Both of the acoustic forces that contribute to microswarm motion Eq. (1 and 2) scale with swarm volume; thus, the larger the swarm, the more prominent the effects of the acoustic forces on it. We examined v_cross_ across vascular flow rates of 4.2 – 16.7 cm/s, a range normally found in human arteries (Fig. 5D). When Piezo transducer 1 was actuated at 242 KHz and 20 V_PP_, cross-stream motion was achieved with velocities up to 3.3 ± 2.02 mm/s. Notably, higher power (20 V_PP_) was needed in the presence of flow, whereas 1 – 5 V_PP_ was sufficient under quiescent flow conditions. Cross-stream allows the microswarms to move towards the wall, where the velocity is minimum due to no slip boundary conditions, in which microrobot manipulation along the wall of the vasculature becomes sustainable (Movie S6).

**Fig. 5.**
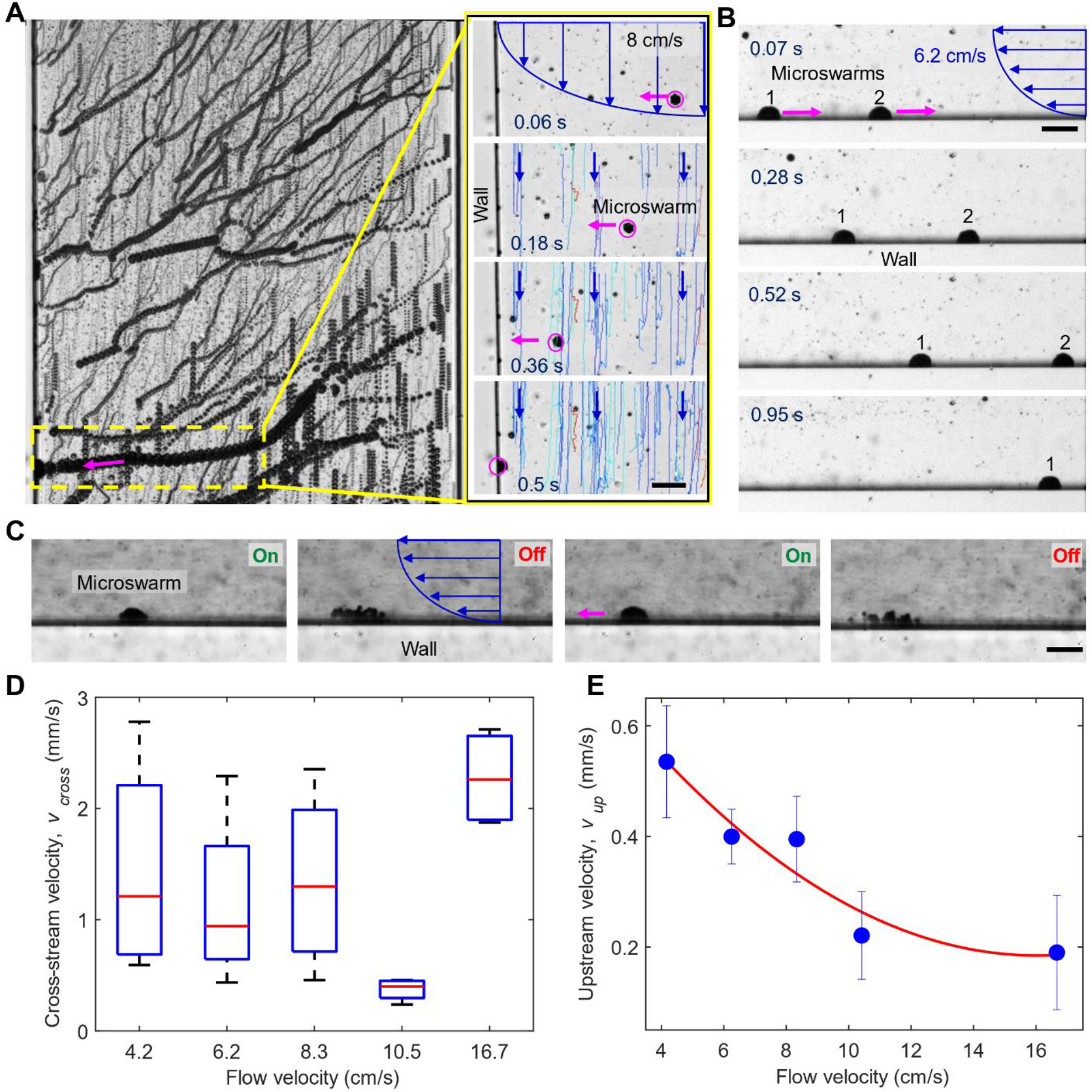
Microswarm manipulation in physiological flow conditions. **(A).** Superimposed time-lapse images demonstrating cross-stream motion of microswarms when acoustics are introduced to the artificial channel. Inset: Image sequence showing flow of microscopic beads inside the channel and downstream tracking in the vessel (blue). A single microswarm, in magenta, was tracked during cross-stream motion towards the wall. **(B).** Upstream motion in microswarms located at the vessel wall. **(C)**. Controlled downstream motion of a cluster located at the wall. When the acoustics are turned off, the swarm starts to disassemble and drift downstream. **(D).** Box plot of microrobot cross-stream velocities under different rates of flow inside the channel. Red line represents median cross-stream velocity analysed from five microrobots. **(E).** Plot of microrobot upstream velocities under different rates of flow inside the channel. Each data point represents the average upstream velocity analysed from five microrobots. Error bars represent standard deviation (s.d.). Scale bar, 50 μm.

Once the microswarms reached the vasculature wall, Piezo transducer 2 was activated at 242 KHz and 20 V_PP_ to exert primary radiation forces parallel to the wall. Most microswarms moving against flow did so only once they were in contact with the wall, where no-slip boundary conditions facilitated upstream motion (Fig. 5B & Movie S7). We characterized this upstream motion in physiologically relevant flow velocities similar to those occurring in human vasculature. As the flow rate increased from 4.2 to 16.7 cm/s, the upstream velocity decreased from 0.53 ± 0.10 mm/s to 0.19 ± 0.10 mm/s, i.e. by ~179%. Higher flow rates resulted in larger drag force on the cluster, thus reducing upstream velocity. Nonetheless, we achieved upstream motion under flow rates of up to 16 cm/s (Fig. 5E), establishing that manipulation inside arteries is feasible under physiological flow rates found in both humans and mice. Importantly, as arteries present the largest flow rates found inside living animals, achieving manipulation in these conditions indicates this approach is robust for *in vivo* use in any type of vessel.

Finally, we also established principles by which to tune swarm disassembly and down-stream motion. Once a microswarm is in contact with a wall, it enters an equilibrium state and remains at the wall until the acoustic field changes. When the acoustic field is turned off, no bubble oscillation takes place, thus bubbles are not attracted either to each other or to the wall; the swarm disassembles immediately and starts following the flow downstream. Note that flow velocity close to the wall is near zero, so that downstream movement proceeds slowly. In addition, vasculature such as our artificial vessel features a laminar flow profile, which is characterized by each component of the flow stream having a straight trajectory, and so all components run parallel to each other. This means that although the microswarm will disassemble slowly and move downstream, its components remain next to the wall while doing so and do not migrate to other parts of the vessel. Thus, as soon as the acoustic field is turned back on, the swarm again self-assembles and presses against the vessel wall (Fig. 5C). With this method for controlled downstream motion, we achieve full control of microswarm motion in a vessel subject to flow.

### The effect of blood microenvironment

When mimicking physiological conditions and investigating the functionality of our system, it is necessary to consider blood viscosity (1.8 times greater than water) and heterogeneity (i.e. components such as red blood cells, platelets, etc.) as the various blood components could impair swarm manipulation. To model the physiological environment and explore the translatability of our previous results obtained in water, we mixed mouse blood with MB-containing saline solution, then used that diluted blood in the artificial vessels into which MBs were injected. We tested two different dilutions and associated flow rates. First, we used a 1:2 ratio of MB saline solution to mouse blood with a flow rate of 0.5 mm/s (Fig. 6A). Acoustic signal was introduced from Piezo transducer 1 at 242 kHz / 5 V and PZT 2 at 242 kHz / 3V. This first preparation enabled efficacious MB tracking during microswarm actuation and manipulation on account of the lower concentration of red blood cells and low flow velocity. Secondly, we employed a 1:6 ratio of MB saline solution to mouse blood with a flow velocity of 1.4 mm/s (Fig. 6B). Acoustic signals were introduced from PZT 1 at 242 kHz / 10 V and PZT 2 at 242 kHz / 3V. This second assessment closely mimicked the physiological environment found inside human or mouse vasculatures. Note that the more concentrated blood preparation damped the acoustic signal and so mandated a larger actuation voltage, i.e. 10 V versus 5 V, for microswarm formation and manipulation.

**Fig. 6.**
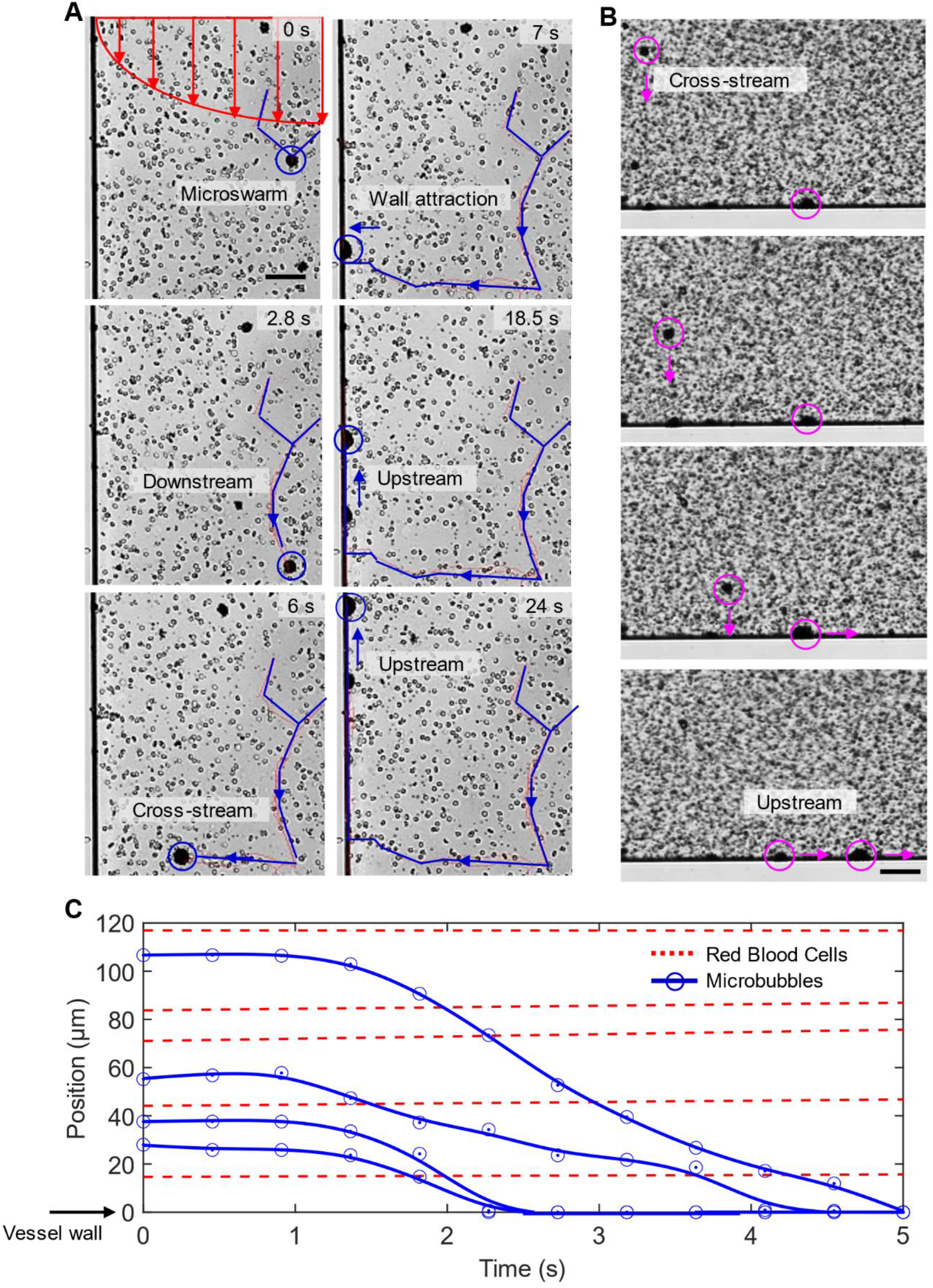
Microswarm manipulation inside blood flow. **(A).** Microswarms are manipulated cross-stream and upstream in the presence of blood. The blood sample used here was less concentrated than *in vivo* mouse blood. One manipulation cycle is shown in which microswarms traverse downstream, cross-stream, and upstream. **(B).** Demonstration of cross-stream and upstream microrobot manipulation in a blood sample with concentration as found in living organisms. **(C).** Comparison of MB and red blood cell behaviours in the presence of both ultrasound and flow. Manual tracking revealed microbubbles to move towards the wall, while RBCs followed a straight downstream trajectory.

During these assessments, microswarm self-assembly was triggered inside a microchannel with an existing blood flow (Movie S7). We observed an initial downstream motion, i.e. the microswarms followed the flow trajectory, followed by cross-stream motion in which the microswarm moved left towards the wall. The cross-stream motion was induced by an acoustic actuator on the right side of the artificial channel (Fig. 6A). Subsequently, we activated an acoustic actuator placed at the bottom, perpendicular to the channel, to move the microswarms upstream while they remained adjacent to the vessel wall. Under both scenarios, cross-stream motion occurred at 104.5 – 29.6 μm/s and upstream motion at 20.7 – 43.6 μm/s. We show in Movie S7 & 8, that not only are cross-stream and upstream manipulation accomplished, but also that the onset of each motion can be triggered at any moment provided the MBs are circulating within the bloodstream. Thus, changing the fluid medium inside the artificial vessel did not impair acoustic manipulation of the MBs or microswarms.

Meanwhile, the trajectories of red blood cells were unaffected (Fig. 6C). That is, the air-filled MBs were selectively activated by the acoustic field, exhibiting a parabolic trajectory towards the wall, while red blood cells strictly demonstrated downstream laminar flow in the vessel. This factor defines how much acoustic force is felt by an object, and one of the key parameters defining it is particle compressibility. As mentioned before, MBs suffer volumetric oscillations when exposed to acoustic waves; the same is true for any compressible object, for example cells. Furthermore, the larger the oscillations, the higher the forces experienced by the object. The acoustic force responsible for particle oscillations can be expressed as(*53*):

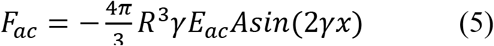

where F_ac_ is the acoustic force, *E_ac_* is the acoustic energy density, γ is the acoustic wave number, and A is a constant factor defined as follows: 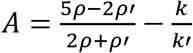, Eq. (6) where *ρ′* is the density of the particle, *k* is the compressibility of the medium, and *k’* is the compressibility of the particle. From Eq. 5 and 6, we derive that the higher the particle compressibility, the more acoustic force is exerted. The gas-filled MBs utilized in our approach are significantly more compressible, than the cellular components present in the bloodstream, i.e. MB isentropic compressibility values(*54*) around 1 versus RBC compressibility of 34.8*10^-12^ (*55*); thus, the MBs can be selectively actuated at acoustic pressures lower than those required to induce oscillation in cells.

### The effect of pulsatile flow

The heart of a human adult at rest executes between 60 and 100 beats per minute (bpm), and the resulting pulsatile flow wave is transmitted along the vasculature network(*56*), with blood flow reaching velocities on the order of cm/s, as mentioned previously. Such pulsatile flow is a feature found mainly in big arteries; as the vasculature branches into smaller vessels, resistance exerted against the flow increases, and the pulsatile wave becomes damped, though it remains a feature even in small arteries(*56*). Accordingly, microswarm functionality in pulsatile flow conditions is an extra consideration that needs to be taken into account for *in vivo* applications. Here, we demonstrated self-assembly and cross-stream motion under a pulse regime of 60 to 180 bpm, i.e., at values within the pulsatility range that occurs in a human body. We implemented pulsatile flow profile inside our artificial channel, as shown in Fig. S3. When applying ultrasound from Piezo transducer 1 followed by Piezo transducer 2, the microswarms performed cross-stream motion at velocities of 0.3 - 1.4 mm/s, and upon reaching the wall subsequently executed upstream movement in the range of 0.5 – 3.5 mm/s, Movie S9.

Notably, the pulsatile flow featured two distinct phases: a low-flow phase with flow velocity between 0 and 1 mm/s, and a high-flow phase with higher velocity between 1 and 3.27 mm/s; these values correspond to manipulation in pulsatile flow of 100 bpm (used for the subsequent analysis). The two differentiated flow phases exerted effects evident in the angled trajectory of cross-stream motion. We observed trajectories in pulsatile flow to be more perpendicular during low-flow phase, with angles of up to 74.9 ± 12.7° in low-flow and around 16.6 ± 9.14° under high-flow (Fig. 7). During the high-flow phase, MBs are subject to both acoustic force (perpendicular to the vessel) and drag force (parallel to flow velocity), leading to a parabolic trajectory. In addition, flow phases affected also net cross-stream velocities; during low-flow, cross-stream velocities were in the range of 1.18 ± 0.46 mm/s and the motion aimed directly towards the wall, while in high-flow the angular motion derived into lower net cross-stream velocities of 0.77 ± 0.31 mm/s. Taken together, we concluded that the cross-stream motion bringing microswarms towards the wall, is influenced but not disrupted by pulsatile flow; microswarms take advantage of low-flow phases between cycles to develop a more effective and directed motion. Once at the wall, microswarms again exerted upstream manipulation on account of low flow values found at boundaries. With the knowledge gained from these results, we can develop a process for controlled manipulation of microrobots in pulsatile conditions.

**Fig. 7.**
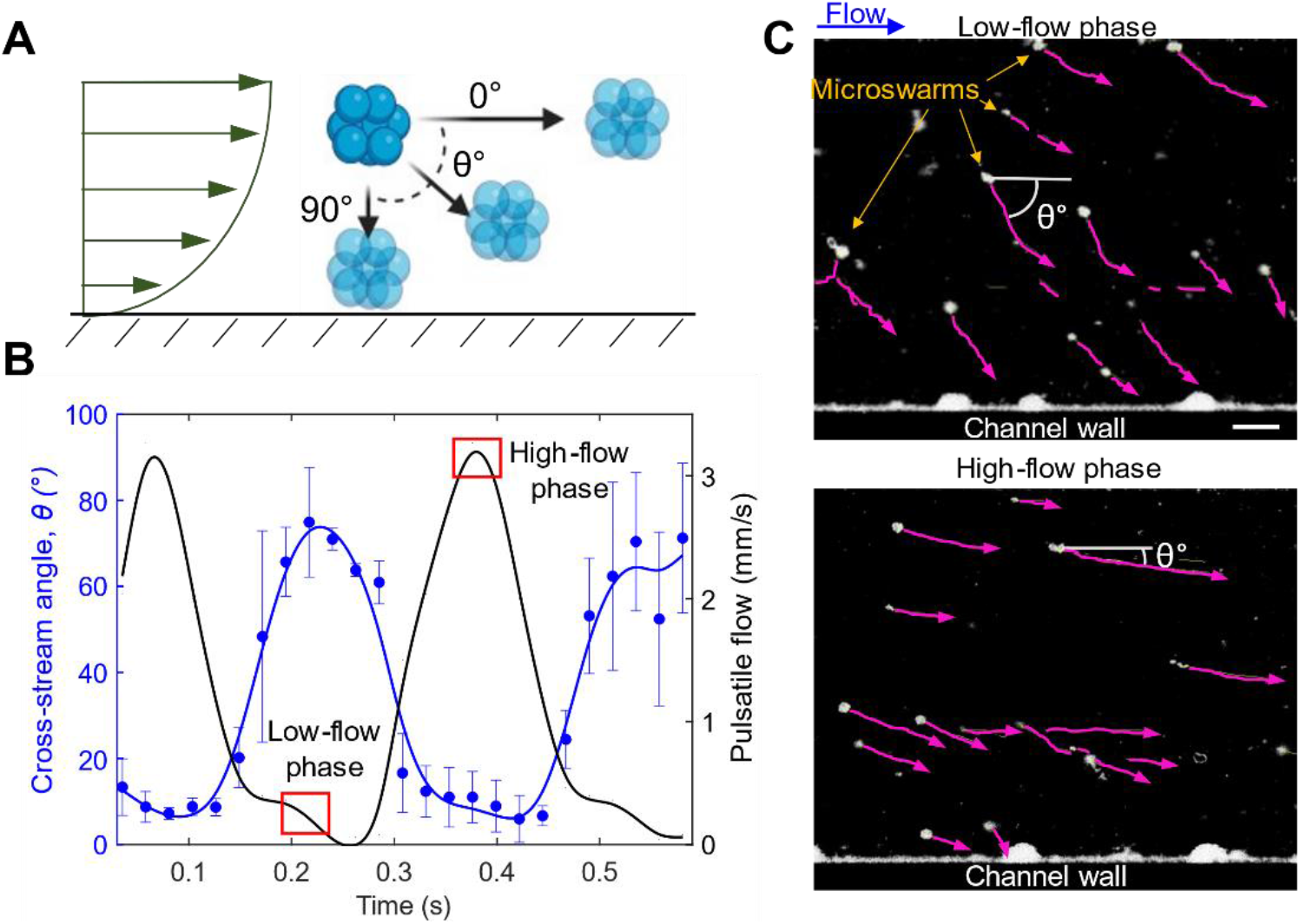
Microswarm manipulation in pulsatile flow conditions. **(A).** Schematic illustrating the concept of cross-stream angle. **(B).** Plot showing the change in cross-stream angle over time during different phases of a 100-bpm pulsatile flow. Each data point represents the average cross-stream angle analysed from five microrobots. Error bars represent standard deviation (s.d.) **(C).** Microswarms were tracked using ImageJ tool TrackMate during their cross-stream motion towards the wall. We demonstrate the difference in cross-stream angles between low-flow and high-flow phases of pulsatile flow.

## Discussion

We introduced a robust and stable approach for microrobot manipulation. The system’s actuation is based in fundamental acoustofluidic physics, where acoustic radiation forces are the principal components controlling microrobot behaviour, and we generate these acoustic forces from only two piezoelectric actuators. Once activated, the microrobots instantaneously begin moving cross-stream towards the minimum flow region located next to the vessel wall. Once the microrobot swarms are stabilized at the boundary, manipulation up-stream becomes less challenging and exhibits highly reproducible results. Notably, the strong and fast response observed in acoustic-activated MBs enables the system to work even in complex environment conditions. Thus, we provide a platform that is coherent, versatile, and meant to be translated to *in vivo* applications.

Our acoustic-driven system is capable of realizing both rapid swarm movement upstream and precise movement in down-stream and cross-stream directions. It merits mention that the observed cross-stream velocities did not entirely align with expected results, as cross-stream motion was expected to depend on flow velocity inside the vessel. Despite this, our system achieved precise manipulation of microswarms even under challenging conditions, i.e. flow rates of up to 16 cm/s. The success of this system in pulsatile flow opens up new possibilities in the world of microrobots, potentially allowing manipulation not only in small capillaries but also in big arteries. Furthermore, in addition to mimicking flow velocities found in human vasculature, we also mimicked the blood microenvironment, including cellular components and pulsatile flow regimes. Thus, we demonstrate great support of the translatability of this system for medical applications. All in all, we provide a detailed characterization of acoustic-mediated microrobot control in terms of assembly speed, swarm size, and manipulation velocities. This characterization will be the foundation for further evaluation of this system in more complex set-ups and finally in *in vivo* scenarios.

Normally, prefabricated robots are limited in their biocompatibility, are difficult and expensive to mass-produce, and require additional surface modifications that can pose challenges for *in vivo* applications. Our self-assembled microbubble-based microswarm systems not only overcome these limitations but also provide additional benefits. In particular, they are contrast agents typically used for ultrasound imaging, and so allow for future real-time tracking and directed navigation by means of *in vivo* imaging. Additionally, a lot of current existing approaches make use of rotational magnetic fields and vortex-shaped acoustic fields, both providing a precise platform for manipulation with high system control; however, blood vessels are lined by endothelial cells, and the creation of vortexes or abnormal flow profiles may induce stresses that disrupt normal endothelial cell function and trigger proinflammatory responses. Such disruption of the blood microenvironment should be minimized in any medical treatment. These swarmbots aim to minimize the shear stress generated during manipulation. In our experiments, no acoustic streaming vortexes were observed around microbubbles or microswarms at power values in the range of 0.1-10 V. At power values >15 V, acoustic streaming that disrupted normal blood flow started to appear; however, those values are greater than what is needed to activate our microrobots. Thus, our microrobots migrate within flow without disrupting it, which helps to minimize their invasiveness. These first results are promising for the development of a minimally invasive microrobot platform. Moreover, since we are using technologies already having clinical approval, these self-assembled microrobots can be fast-tracked into translational medicine. Our technology will have significant impact as a foundation for advancements in brain research and vasculature biology, as well as in furthering our understanding of diseases and the development of relevant treatments. Future work will focus on the implementation of the system in more complex channel-network geometries and 3D designs, and to prove the validity of the system for *in vivo*_=_studies_≐_

## Materials and Methods

### Acoustofluidic device and experimental set-up

The microfluidic design used for the experiments was composed of a single channel, 1.5 mm length and 400 × 30 μm cross-section. We used soft lithography methodology to finally transfer the channel design to a PDMS slab. Details on soft lithography steps can be found in the Supplementary information. We attached two PDMS blocks together resting on a glass slide, and the microfluidic channel lying in between. We employed plasma treatment for 10 seconds to their inner surfaces, we attached them together and applied heat treatment at 85°C for 1 hour. A Steminc Piezo Ceraminc Plate 7×8×0.2mm, Wire Lead, 240 KHz resonant frequency, was used as acoustic transducer. We placed the piezo electric acoustic actuator bonded next to the microfluidic chamber by using 2-component epoxy glue, ensuring a good coupling between the piezo transducer and the PDMS wall. The acoustic transducer was placed so that there was no contact between it and the glass slide. We actuated the piezo transducer connecting it to an Arbitrary Function Generator, Tektronix. We used computer-controlled programmable syringe pumps, connected via a silicon tube to the channel inlet, to regulate the liquid flow within. A 5ml syringe was used to gather the pumped fluid. The whole setup was mounted on an inverted microscope (Zeiss 200m/Nikon Eclipse Ti-U), and images were captured using either an AxioCAM MrM, Photometrics HQ2 high-sensitive camera or a Chronos high-speed camera.

### Microbubble contrast agents

The MBs used for microswarm assembly were commercially purchased Bracco Sonovue imaging contrast agent. Sonovue contrast agent comes inside a ‘fabrication kit’ which includes: a glass vial containing 25mg of lyophilised sulphur hexafluoride lipid-type A powder, one pre-filled syringe containing 5 ml sodium chloride (saline) solution and one Mini-Spike transfer system. A microsphere dispersion is created after injecting saline solution into the glass vial and a gentle shaking. We made different MB concentration dilutions using the saline solution provided; so that different MB concentrations could be tested. We injected the MB solution with a pipette into the microfluidic channel and also into the silicon tube connected to the syringe pump. Thus, even under the presence of a flow, freshly new bubbles entered into the channel. The self-assembly of the microswarms was recorded by a camera connected to an inverted microscope, as mentioned above, and analysed using MATLAB and ImageJ.

### Preparation of blood

Mice blood samples were kept stored in ice while performing the experiments. For the experiments, blood was mixed with MB saline solution. The blood/MB mix was introduced with a pipette inside the microfluidic channel and inside the syringe pump silicon tube. The self-assembly of the microswarms in this environment was again recorded by a camera connected to an inverted microscope and analysed using MATLAB and ImageJ.

## Author Contributions

D.A. conceived and supervised the project. A.C.F and T.K. performed all the experiments and performed data analysis with feedback from D.A. All authors contributed to the experimental design, scientific presentation, and discussion. A.C.F and D.A. wrote the manuscript.

## Data and materials availability

All data are available in the main text or the supplementary materials.

## References

1. S. Palagi, P. Fischer, Bioinspired microrobots. Nat Rev Mater. 3, 113–124 (2018).

2. O. Aydin, X. Zhang, S. Nuethong, G. J. Pagan-Diaz, R. Bashir, M. Gazzola, M. T. A. Saif, Neuromuscular actuation of biohybrid motile bots. PNAS. 116, 19841–19847 (2019).

3. O. Aydin, A. P. Passaro, M. Elhebeary, G. J. Pagan-Diaz, A. Fan, S. Nuethong, R. Bashir, S. L. Stice, M. T. A. Saif, Development of 3D neuromuscular bioactuators. APL Bioengineering. 4, 016107 (2020).

4. K. Belharet, D. Folio, A. Ferreira, Simulation and planning of a magnetically actuated microrobot navigating in the arteries. IEEE Trans Biomed Eng. 60, 994–1001 (2013).

5. X. Ding, S.-C. S. Lin, B. Kiraly, H. Yue, S. Li, I.-K. Chiang, J. Shi, S. J. Benkovic, T. J. Huang, On-chip manipulation of single microparticles, cells, and organisms using surface acoustic waves. PNAS. 109, 11105–11109 (2012).

6. Z. Li, M. T. A. Saif, Mechanics of Biohybrid Valveless Pump-Bot. Journal of Applied Mechanics. 88 (2021).

7. J. Llacer-Wintle, A. Rivas-Dapena, X.-Z. Chen, E. Pellicer, B. J. Nelson, J. Puigmartí-Luis, S. Pané, Biodegradable Small-Scale Swimmers for Biomedical Applications. Advanced Materials. n/a, 2102049 (2021).

8. J. Shi, X. Mao, D. Ahmed, A. Colletti, T. Jun Huang, Focusing microparticles in a microfluidic channel with standing surface acoustic waves (SSAW). Lab on a Chip. 8, 221–223 (2008).

9. S. Palagi, D. P. Singh, P. Fischer, Light-Controlled Micromotors and Soft Microrobots. Advanced Optical Materials. 7, 1900370 (2019).

10. Z. Cenev, P. A. D. Harischandra, S. Nurmi, M. Latikka, V. Hynninen, R. H. A. Ras, J. V. I. Timonen, Q. Zhou, Ferrofluidic Manipulator: Automatic Manipulation of Nonmagnetic Microparticles at the Air–Ferrofluid Interface. IEEE/ASME Transactions on Mechatronics. 26, 1932–1940 (2021).

11. X. Dong, G. Z. Lum, W. Hu, R. Zhang, Z. Ren, P. R. Onck, M. Sitti, Bioinspired cilia arrays with programmable nonreciprocal motion and metachronal coordination. Science Advances. 6, eabc9323 (2020).

12. Y. Alapan, B. Yigit, O. Beker, A. F. Demirörs, M. Sitti, Shape-encoded dynamic assembly of mobile micromachines. Nat. Mater. 18, 1244–1251 (2019).

13. D. Folio, A. Ferreira, Two-Dimensional Robust Magnetic Resonance Navigation of a Ferromagnetic Microrobot Using Pareto Optimality. IEEE Transactions on Robotics. 33, 583–593 (2017).

14. I. S. M. Khalil, H. Abass, M. Shoukry, A. Klingner, R. M. El-Nashar, M. Serry, S. Misra, Robust and Optimal Control of Magnetic Microparticles inside Fluidic Channels with Time-Varying Flow Rates. International Journal of Advanced Robotic Systems. 13, 123 (2016).

15. K. Belharet, D. Folio, A. Ferreira, in 2012 IEEE/RSJ International Conference on Intelligent Robots and Systems (2012), pp. 2559–2564.

16. H. Xu, M. Medina-Sánchez, M. F. Maitz, C. Werner, O. G. Schmidt, Sperm Micromotors for Cargo Delivery through Flowing Blood. ACS Nano. 14, 2982–2993 (2020).

17. D. Xu, J. Hu, X. Pan, S. Sánchez, X. Yan, X. Ma, Enzyme-Powered Liquid Metal Nanobots Endowed with Multiple Biomedical Functions. ACS Nano. 15, 11543–11554 (2021).

18. A. C. Hortelao, C. Simó, M. Guix, S. Guallar-Garrido, E. Julián, D. Vilela, L. Rejc, P. Ramos-Cabrer, U. Cossío, V. Gómez-Vallejo, T. Patiño, J. Llop, S. Sánchez, Swarming behavior and in vivo monitoring of enzymatic nanomotors within the bladder. Science Robotics. 6, eabd2823 (2021).

19. D. Vilela, N. Blanco-Cabra, A. Eguskiza, A. C. Hortelao, E. Torrents, S. Sanchez, Drug-Free Enzyme-Based Bactericidal Nanomotors against Pathogenic Bacteria. ACS Appl. Mater. Interfaces. 13, 14964–14973 (2021).

20. V. Sridhar, F. Podjaski, J. Kröger, A. Jiménez-Solano, B.-W. Park, B. V. Lotsch, M. Sitti, Carbon nitride-based light-driven microswimmers with intrinsic photocharging ability. PNAS. 117, 24748–24756 (2020).

21. C. Meredith, A. Castonguay, Y.-J. Chiu, A. M. Brooks, P. Moerman, P. Torab, P. K. Wong, A. Sen, D. Velegol, L. Zarzar, Chemical Design of Self-Propelled Janus Droplets. Chem. Rev. (2021).

22. A. Somasundar, A. Sen, Chemically Propelled Nano and Micromotors in the Body: Quo Vadis? Small. 17, 2007102 (2021).

23. J. Katuri, W. E. Uspal, J. Simmchen, A. Miguel-López, S. Sánchez, Cross-stream migration of active particles. Science Advances. 4, eaao1755.

24. D. Ahmed, T. Baasch, N. Blondel, N. Läubli, J. Dual, B. J. Nelson, Neutrophil-inspired propulsion in a combined acoustic and magnetic field. Nature Communications. 8, 1–8 (2017).

25. D. Ahmed, A. Sukhov, D. Hauri, D. Rodrigue, G. Maranta, J. Harting, B. J. Nelson, Bioinspired acousto-magnetic microswarm robots with upstream motility. Nat Mach Intell. 3, 116–124 (2021).

26. Y. Alapan, U. Bozuyuk, P. Erkoc, A. C. Karacakol, M. Sitti, Multifunctional surface microrollers for targeted cargo delivery in physiological blood flow. Science Robotics (2020), doi:10.1126/scirobotics.aba5726.

27. Q. Wang, K. F. Chan, K. Schweizer, X. Du, D. Jin, S. C. H. Yu, B. J. Nelson, L. Zhang, Ultrasound Doppler-guided real-time navigation of a magnetic microswarm for active endovascular delivery. Science Advances. 7, eabe5914 (2021).

28. S. R. Goudu, I. C. Yasa, X. Hu, H. Ceylan, W. Hu, M. Sitti, Biodegradable Untethered Magnetic Hydrogel Milli-Grippers. Advanced Functional Materials. 30, 2004975 (2020).

29. P. Fischer, D. Sanz-Hernández, R. Streubel, A. Fernández-Pacheco, Launching a new dimension with 3D magnetic nanostructures. APL Materials. 8, 010701 (2020).

30. D. Sanz-Hernández, A. Hierro-Rodriguez, C. Donnelly, J. Pablo-Navarro, A. Sorrentino, E. Pereiro, C. Magén, S. McVitie, J. M. de Teresa, S. Ferrer, P. Fischer, A. Fernández-Pacheco, Artificial Double-Helix for Geometrical Control of Magnetic Chirality. ACS Nano. 14, 8084–8092 (2020).

31. X. Liu, N. Kent, A. Ceballos, R. Streubel, Y. Jiang, Y. Chai, P. Y. Kim, J. Forth, F. Hellman, S. Shi, D. Wang, B. A. Helms, P. D. Ashby, P. Fischer, T. P. Russell, Reconfigurable ferromagnetic liquid droplets. Science. 365, 264–267 (2019).

32. H. Zhou, C. C. Mayorga-Martinez, S. Pané, L. Zhang, M. Pumera, Magnetically Driven Micro and Nanorobots. Chem. Rev. 121, 4999–5041 (2021).

33. H. Zhou, C. C. Mayorga-Martinez, M. Pumera, Microplastic Removal and Degradation by Mussel-Inspired Adhesive Magnetic/Enzymatic Microrobots. Small Methods. 5, 2100230 (2021).

34. B. Wang, K. F. Chan, K. Yuan, Q. Wang, X. Xia, L. Yang, H. Ko, Y.-X. J. Wang, J. J. Y. Sung, P. W. Y. Chiu, L. Zhang, Endoscopy-assisted magnetic navigation of biohybrid soft microrobots with rapid endoluminal delivery and imaging. Science Robotics. 6, eabd2813 (2021).

35. S. Pane, V. Iacovacci, E. Sinibaldi, A. Menciassi, Real-time imaging and tracking of microrobots in tissues using ultrasound phase analysis. Appl. Phys. Lett. 118, 014102 (2021).

36. A. Aziz, S. Pane, V. Iacovacci, N. Koukourakis, J. Czarske, A. Menciassi, M. Medina-Sánchez, O. G. Schmidt, Medical Imaging of Microrobots: Toward In Vivo Applications. ACS Nano. 14, 10865–10893 (2020).

37. W. Wang, L. A. Castro, M. Hoyos, T. E. Mallouk, Autonomous motion of metallic microrods propelled by ultrasound. ACS nano. 6, 6122–6132 (2012).

38. J. McNeill, N. Sinai, J. Wang, V. Oliver, E. Lauga, F. Nadal, T. E. Mallouk, Purely viscous acoustic propulsion of bimetallic rods. Phys. Rev. Fluids. 6, L092201 (2021).

39. J. Li, T. Li, T. Xu, M. Kiristi, W. Liu, Z. Wu, J. Wang, Magneto-acoustic hybrid nanomotor. Nano Letters. 15, 4814–4821 (2015).

40. W. Wang, S. Li, L. Mair, S. Ahmed, T. J. Huang, T. E. Mallouk, Acoustic Propulsion of Nanorod Motors Inside Living Cells. Angewandte Chemie. 126, 3265–3268 (2014).

41. D. Ahmed, T. Baasch, B. Jang, S. Pane, J. Dual, B. J. Nelson, Artificial Swimmers Propelled by Acoustically Activated Flagella. Nano Letters. 16, 4968–4974 (2016).

42. A. Aghakhani, O. Yasa, P. Wrede, M. Sitti, Acoustically powered surface-slipping mobile microrobots. PNAS. 117, 3469–3477 (2020).

43. D. Ahmed, C. Dillinger, A. Hong, B. J. Nelson, Artificial Acousto-Magnetic Soft Microswimmers. Advanced Materials Technologies. 2, 1700050 (2017).

44. N. F. Läubli, M. S. Gerlt, A. Wüthrich, R. T. M. Lewis, N. Shamsudhin, U. Kutay, D. Ahmed, J. Dual, B. J. Nelson, Embedded Microbubbles for Acoustic Manipulation of Single Cells and Microfluidic Applications. Anal. Chem. 93, 9760–9770 (2021).

45. P. Dayton, A. Goode, K. Morgan, S. Klibanov, G. Brandenburger, K. Ferrara, Action of microbubbles when insonified: experimental evidence. 1996 IEEE Ultrasonics Symposium. Proceedings (1996).

46. K. Masuda, R. Koda, N. Watarai, N. Shigehara, T. Ito, T. Kakimoto, Y. Miyamoto, S. Enosawa, T. Chiba, Experimental study of active control of microbubbles in blood flow by forming their aggregations. 2011 IEEE International Ultrasonics Symposium (2011).

47. P. Dayton, A. Klibanov, G. Brandenburger, K. Ferrara, Acoustic radiation force in vivo: a mechanism to assist targeting of microbubbles. Ultrasound Med Biol. 25, 1195–1201 (1999).

48. J. Li, C. Shen, T. J. Huang, S. A. Cummer, Acoustic tweezer with complex boundary-free trapping and transport channel controlled by shadow waveguides. Science Advances. 7, eabi5502.

49. P. Kang, Z. Tian, S. Yang, W. Yu, H. Zhu, H. Bachman, S. Zhao, P. Zhang, Z. Wang, R. Zhong, T. J. Huang, Acoustic tweezers based on circular, slanted-finger interdigital transducers for dynamic manipulation of micro-objects. Lab Chip. 20, 987–994 (2020).

50. Y. Yoshiki, O. Yoshiyuki, I. Masato, M. Nobuyuki, Effects of Bjerknes Forces on Gas-Filled Microbubble Trapping by Ultrasonic Waves. Japanese Journal of Applied Physics. 40, 3852–3855 (2001).

51. M. Klarhöfer, B. Csapo, C. Balassy, J. C. Szeles, E. Moser, High-resolution blood flow velocity measurements in the human finger. Magn Reson Med. 45, 716–719 (2001).

52. I. T. Gabe, J. H. Gault, J. Ross, D. T. Mason, C. J. Mills, J. P. Schillingford, E. Braunwald, Measurement of Instantaneous Blood Flow Velocity and Pressure in Conscious Man with a Catheter-Tip Velocity Probe. Circulation. 40, 603–614 (1969).

53. K. Yasuda, S. Umemura, K. Takeda, Concentration and Fractionation of Small Particles in Liquid by Ultrasound. Jpn. J. Appl. Phys. 34, 2715 (1995).

54. P. Marmottant, S. van der Meer, M. Emmer, M. Versluis, N. de Jong, S. Hilgenfeldt, D. Lohse, A model for large amplitude oscillations of coated bubbles accounting for buckling and rupture. The Journal of the Acoustical Society of America. 118, 3499–3505 (2005).

55. K. K. Shung, B. A. Krisko, J. O. Ballard, Acoustic measurement of erythrocyte compressibility. The Journal of the Acoustical Society of America. 72, 1364–1367 (1982).

56. Y. Huo, G. S. Kassab, Pulsatile blood flow in the entire coronary arterial tree: theory and experiment. American Journal of Physiology-Heart and Circulatory Physiology. 291, H1074–H1087 (2006).

