## Supplementary information for "Navigation of Ultrasound-controlled Swarmbots under Physiological Flow Conditions"

Supplementary Materials for

### Ultrasound-controlled Swarmbots Capable of Navigating under Physiological Flow Conditions

Alexia Del Campo Fonseca, Tobias Kohler, and Daniel Ahmed*

Acoustic Robotics Systems Lab, Department of Mechanical and Process Engineering, ETH Zurich, Switzerland

**Contents:**

Figs. S1 to S3

Note S1

Video legends

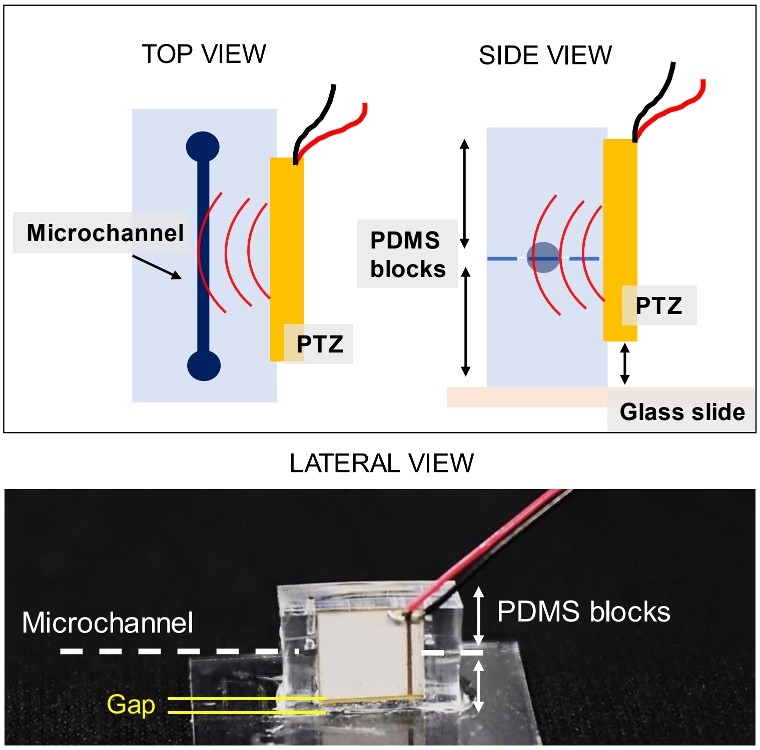


**Supplementary Fig. 1. The experimental setup**. We bonded two PDMS blocks together resting on a glass slide, with a microfluidic channel lying in between. The acoustic actuator (Piezo transducer) was bonded to the lateral surface without touching the glass slide below. A gap was left between the transducer and the glass slide.

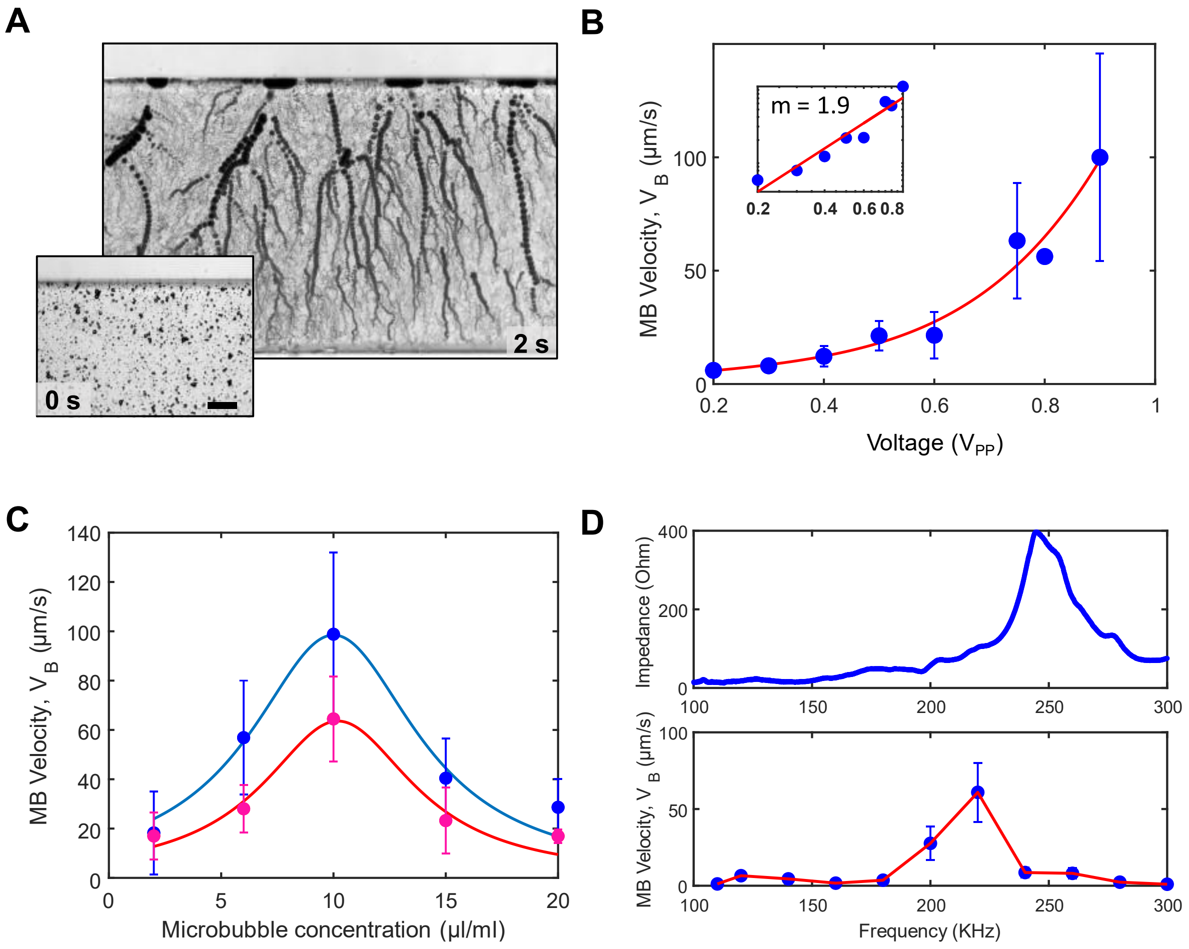


**Supplementary Fig. 2. Characterization of microswarm self-assembly. A.** Superimposed time lapse images showing microswarm self-assembly and migration to the vessel wall. Inset: Initial situation, singe microbubbles dispersed in saline solution. **B**. Plot analysing single microbubble velocities against voltage. Velocities analysed are baseline velocities, $v_B$. **C.** Plot analysing MB velocity versus MB concentration. Blue: single bubble velocity. Magenta: Swarm velocity. **D.** Top: Acoustic impedance of the piezo transducer, when coupled to the experimental setup. Bottom: Plot analysing single MB velocity versus frequency of the acoustic signal.

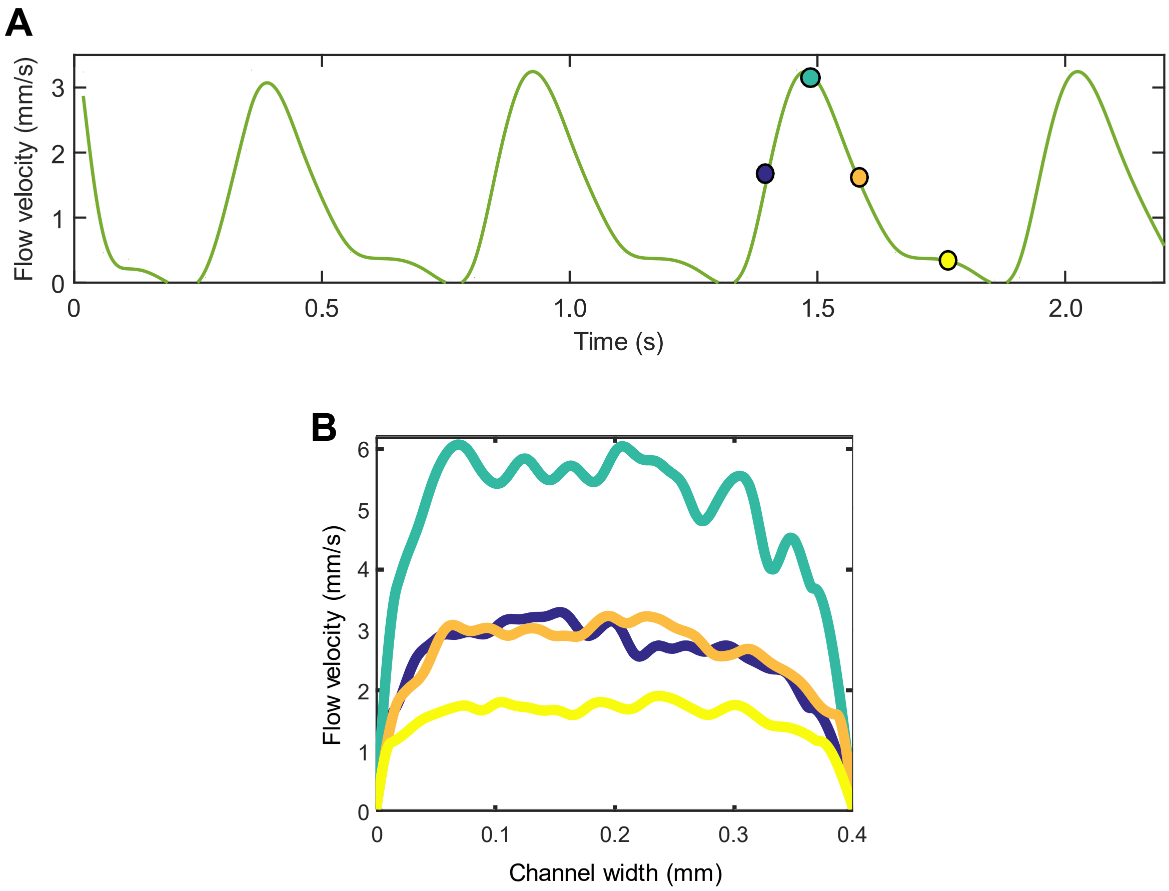


**Supplementary Fig. 3 . Characterization of pulsatile flow. A.** Flow velocity versus time analysed from the pulsatile flow induced in our channels, at 100bpm. **B.** Plot showing the flow profile present inside the channel at different time points of the pulsatile flow shown in ‘a’.

**Supplementary note 1. Soft lithography**

We used a 2D computer-aided design and drafting software, ‘Layout Editor’, to create a draft of the microfluidic channel shapes and geometries. Layout Editor software enabled compatibility with the lithography tool which transferred the design to a photomask substrate, with a microscale resolution. The final design used for the experiments was composed of a single channel, 1.5 mm length and 400 x 30 µm cross-section. We used soft lithography methodology to transfer the channel design from the photomask to a PDMS slab. First, we copied the designs into a silicon substrate via photolithography. For this, we coated a silicon surface with a photoresist and we placed on top of it, the photomask fabricated with our design. Subsequently, light was exposed, triggering a chemical change in the uncovered photoresist which makes it resistant to developer medium. Thus, the device was treated with a developer compound that dissolves the regions of photoresist which were not subjected to light. The device was later etched, transferring the design pattern from the photoresist to the silicon substrate. In the end, the photoresist was no needed any longer, so it was stripped from the wafer.

We used this silicon wafer as mold to copy our channel design into PDMS polymer, via mold replica. We vapor coated the silicon mold with hydrophobic silane solution to enable PDMS later detach from the mold. Subsequently, we mixed silicon elastomer base and curing agent at 10:1 ratio and we casted it onto the silicon mold. We degassed the still uncured PDMS on the silicon mold inside a vacuum chamber, to remove any air microbubbles that would interfere with travelling acoustic signals. Afterwards, we cured the polymer using heat treatment at 85°C for 2 hours. We gently removed the cured PDMS from the silicon mold, and we punched the inlets and the outlets with a 1mm diameter punch.

#### Video Legends

**Supplementary Video 1: Self-assembly of microbubbles into spheres**

(3.26 MB)

The video shows the assembly of microbubbles into a spherical swarm. We applied acoustic excitation at 242 kHz and 1.6 V peak-to-peak. The video was captured at 4434 frames per second (fps) and played at 50 fps.

**Supplementary Video 2: Microswarm self-assembly and navigation to the wall**

(20.6 MB)

The video corresponds to Fig. 2a. The video shows the assembly of microbubbles and simultaneous navigation towards the bottom channel wall. We used TrackMate from Fiji to track single microbubbles. We applied acoustic excitation at 242 kHz and 1.2 V peak-to-peak. The video was captured at 4434 fps and played at 1000 fps.

**Supplementary Video 3: Migration and acceleration of microswarms when approaching the wall**

(4.23 MB)

The video corresponds to Fig. 2b. The video shows the navigation and acceleration of a microswarm towards the channel wall. We applied acoustic excitation at 242 kHz and 3 V peak-to-peak. The video was captured at 4434 fps and played at 250 fps.

**Supplementary Video 4: Microswarm navigation along a wall in a stationary fluid**

(5.6 MB)

The video corresponds to Fig. 3. The video shows the navigation of a microswarm along a wall in a stationary fluid. We applied acoustic excitation at 242 kHz and 5 V peak-to-peak. The video was captured at 8.8 fps and played in real-time.

**Supplementary Video 5: Microswarms self-assembly and migration to the wall in the physiological flow**

(30.22 MB)

The video shows the self-assembly and navigation of microswarms under a flow of 3cm/s. We applied acoustic excitation at 242 kHz and 10 V peak-to-peak. The video was captured at 8.8 fps and played in real-time.

**Supplementary Video 6: Microswarm navigates cross-stream in flow conditions**

(29.35 MB)

The video corresponds to Fig. 5a. The video shows the cross-stream navigation of microswarms under flow conditions. Microbeads flowing in the background were tracked in blue. We applied acoustic excitation at 242 kHz and 20 V peak-to-peak. The video was captured at 1519 fps and played at 50fps.

**Supplementary Video 7: Microswarm navigates upstream in mice blood**

(15.22 MB)

The video corresponds to Fig. 6a. This video shows upstream navigation of microswarms in the presence of blood. We applied acoustic excitation from Piezo transducer 1 at 242 kHz and 5 V peak-to-peak and Piezo transducer 2 at 242 kHz and 3 V peak-to-peak. The video was captured at 8.8 fps and played at real time.

**Supplementary Video 8: Microswarm migration cross-stream and upstream in blood**

(12.3 MB)

The video shows cross-stream and upstream navigation of microswarms in the presence of blood. We applied acoustic excitation from Piezo transducer 1 at 242 kHz and 5 V peak-to-peak and Piezo transducer 2 at 242 kHz and 3 V peak-to-peak. The video was captured at 8.8 fps and played in real-time.

**Supplementary Video 9: Microswarm manipulation under pulsatile flow conditions**

(19.44 MB)

The video shows microswarms navigation in pulsatile flow. We applied acoustic excitation at 242 kHz and 20 V peak-to-peak. The video was captured at 1069 fps and played at 50 fps.
